# Neural signatures of stress susceptibility and resilience in the amygdala-hippocampal network

**DOI:** 10.1101/2023.10.23.563652

**Authors:** Frances Xia, Valeria Fascianelli, Nina Vishwakarma, Frances Grace Ghinger, Stefano Fusi, Mazen A Kheirbek

## Abstract

The neural dynamics that underlie divergent anhedonic responses to stress remain unclear. Here, we identified neuronal dynamics in an amygdala-hippocampal circuit that distinguish stress resilience and susceptibility. In a reward-choice task, basolateral amygdala (BLA) activity in resilient mice showed enhanced discrimination of upcoming reward choices. In contrast, a rumination-like signature emerged in the BLA of susceptible mice; a linear decoder could classify the intention to switch or stay on a previously chosen reward. Spontaneous activity in the BLA of susceptible mice was higher dimensional than controls, reflecting the exploration of a larger number of distinct neural states. Manipulation of vCA1-BLA inputs rescued dysfunctional neural dynamics and anhedonia in susceptible mice, suggesting that targeting this pathway can enhance BLA circuit function and ameliorate of depression-related behaviors.

**One-Sentence Summary:** Identification and rescue of dysfunctional vCA1-BLA population dynamics and behavior in stress-susceptible mice.

## Main Text

Organisms adapt their behavior to increase chances of obtaining high-value rewards. These drives for rewards are particularly influenced by changes in emotional states. Anhedonia, a core feature of depression, is characterized by a reduced ability to experience pleasure (*1*), leading to diminished reward responsiveness, learning, and valuation (*2–5*). In rodent models, while some animals show resilience to chronic stress, a subset of stress-susceptible mice display high levels of social avoidance and reduced preference for high-value rewards (*5–7*). The underlying neural dynamics that may produce these behavioral differences in resilient and susceptible individuals remains a significant and unanswered question.

The reciprocally connected amygdala (AMY) and ventral hippocampus (vHPC) are crucial nodes in an extended network responsible for generating emotional and motivated behavior (*8–37*). In addition to its role in threat detection and anxiety-related behavior, the basolateral amygdala (BLA) generates outcome-specific reward representations to guide decision-making (*38–45*). Ventral CA1 (vCA1) has also been shown to encode reward-predictive stimuli and drive reward-related approach behaviors (*46–50*). However, how these reward-related functions of vCA1 and BLA are impacted by changes in emotional state remain unclear. While stimulus-evoked responses in single neurons in the BLA and vCA1 have been well-studied, we know significantly less about how the reward-related internal states of the animal are represented in the BLA and vCA1 and how they drive choice-related behavior. Moreover, how these reward-related dynamics, or spontaneous patterns of activity, in the BLA or vCA1 may differ in mice susceptible or resilient to chronic stress, and how targeted interventions in this circuit may reduce susceptible phenotypes remain unknown. As the internal states can be detected and studied only by looking at the correlations between the neural activities of multiple neurons, it is essential to record simultaneously from a large number of neurons, and to analyze the activity at the population level. Moreover, compared to single neuron responses, population activity can more accurately reveal the dynamics of internal states. Here, we conducted high-density Neuropixels recordings (*51*) in vCA1 and BLA and used population decoders to analyze the reward-related and spontaneous dynamics of populations of neurons to identify distinctive neural signatures of susceptibility and resilience to chronic stress. Then, we developed a novel circuit-specific modulation approach to rescue aberrant BLA population dynamics and associated anhedonic behavior in stress-susceptible mice.

## Results

### Distinct behavioral signatures of resilient and susceptible mice following CSDS

We performed high-density single unit electrophysiology using Neuropixels probes implanted in BLA and vCA1 in control mice and those subjected to chronic social defeat stress (CSDS) (Fig. 1A-E). We recorded activity both under a stimulus-free pre-task condition and as mice performed a novel head-fixed sucrose preference test (SPT). CSDS produced mice with varying degrees of sucrose preference and social interaction scores that were highly correlated (Fig. 1F). We used these measures to perform unsupervised K-means clustering to classify mice as resilient or susceptible (Fig. 1G). The susceptible mice identified using this classification showed lower lick rates during sucrose consumption, as well as reduced lick rate discrimination between sucrose and water rewards, behavioral subcomponents that reflect anhedonia (*5*) (Fig. 1H-I and S1A-B).

**Fig. 1.**
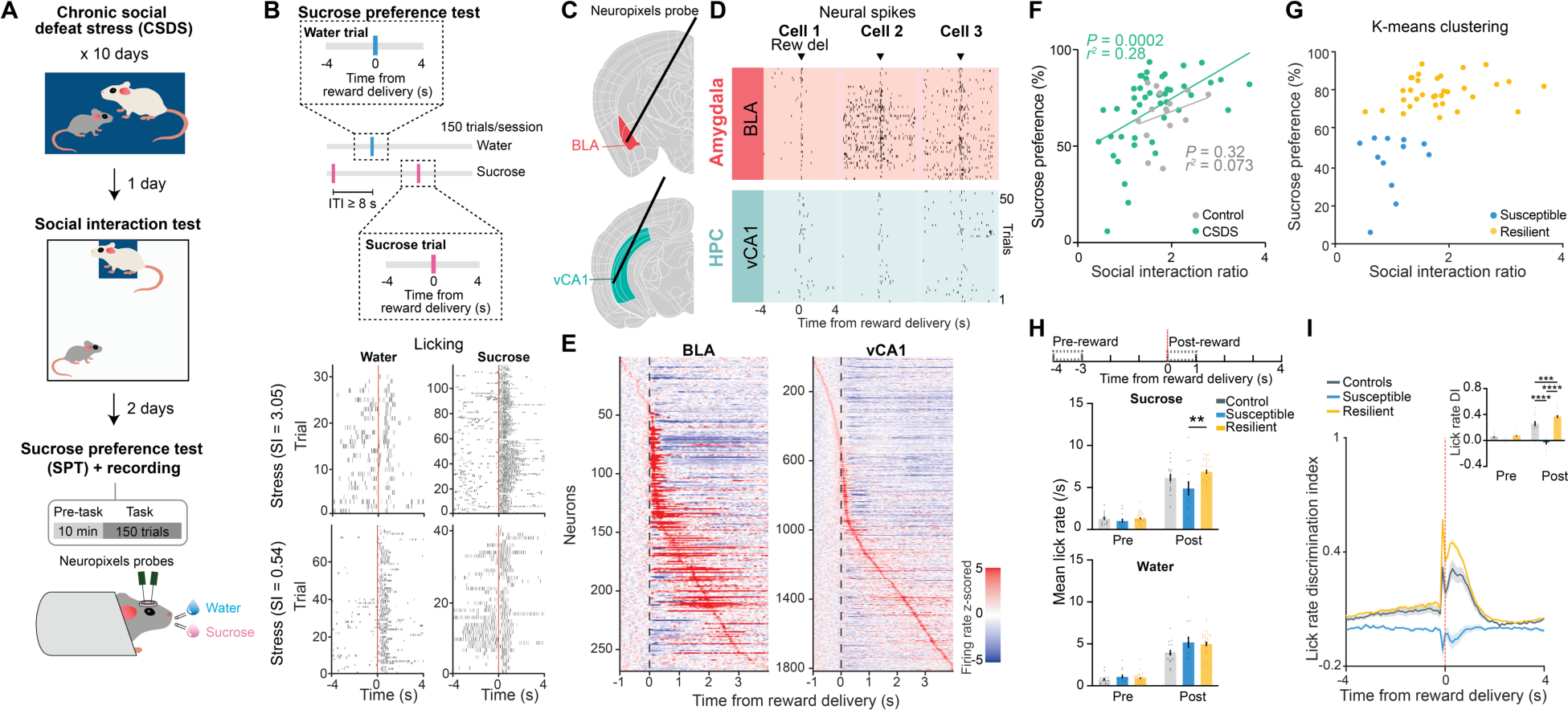
Distinct behavioral signatures of resilient and susceptible mice following CSDS. (**A**) Schematic of SPT and Neuropixels recording protocol following CSDS. (**B**) Schematic of SPT protocol. Lick rasters were from two representative mice with different sucrose preferences. (**C**) Neuropixels probes were targeted to BLA and vCA1. (**D**) Example BLA and vCA1 spike rasters during task. (**E**) Example peristimulus time histogram of BLA and vCA1 neurons around the time of reward delivery (time 0). (**F**) CSDS, but not control, mice showed a significant correlation between sucrose preference and social interaction ratio. (**G**) Unsupervised K-means clustering revealed 2 distinct subgroups of CSDS mice. (**H**) Susceptible mice showed reduced sucrose lick rate during Post-reward period compared to resilient mice (RM-ANOVA, group x time interaction: *F*_2,57_ = 5.63, *P* = 0.0059). (**I**) Susceptible mice showed reduced lick rate discrimination index compared to control and resilient mice (RM-ANOVA, group x time interaction: *F*_2,57_ = 48.47, *P* < 0.0001). Data are mean ± SEM. # Significantly different from chance; * *P* < 0.05; ** *P* < 0.01.

### Enhanced discrimination of reward choice in stress resilient mice

As we observed robust sucrose-seeking behaviors in resilient mice as compared to susceptible mice, we looked for specific adaptations in neural representations of reward-related information. Mice were allowed to freely choose water or sucrose and indicated their choice by licking a spout to trigger reward delivery (Fig. 2A). We defined a trial type (sucrose or water) using a time window around reward delivery (4s Pre-to 4s Post-reward), as it allowed us to assess neural activity patterns before the mice behaviorally made their choices and after they consumed the reward. When quantifying reward-choice-selectivity of each recorded BLA and vCA1 neuron, we found that resilient mice had the highest proportion of reward-choice-selective neurons in the BLA during both reward consumption (Post-reward) and in the seconds before reward delivery (Pre-reward) compared to susceptible and control mice (Fig. 2B-C). This difference in stress groups was specific for BLA as, in vCA1, we found that stress increased the proportion of reward-choice-selective neurons in all mice. To investigate differences between groups at the population level, we trained linear classifiers to discriminate trial types, balancing the number of current and past rewards for each trial type (Fig. 2D, see Methods). When analyzing Pre-reward time bins, we again found that in resilient mice, upcoming reward choice could be decoded from neural activity in BLA better than chance and better than from neurons in control and susceptible mice (Fig. 2E-F). This effect in resilient mice was more robust after a previous low-value water trial than a high-value sucrose trial, which may drive resilient mice to preferentially seek out sucrose rewards (Fig. S2A-B). We observed a similar phenomenon after reward consumption in the BLA (Post-reward), while in vCA1, there existed a general effect of stress. These results suggest that BLA neurons in resilient mice showed enhanced discrimination of reward identities both before and during reward consumption.

**Fig. 2.**
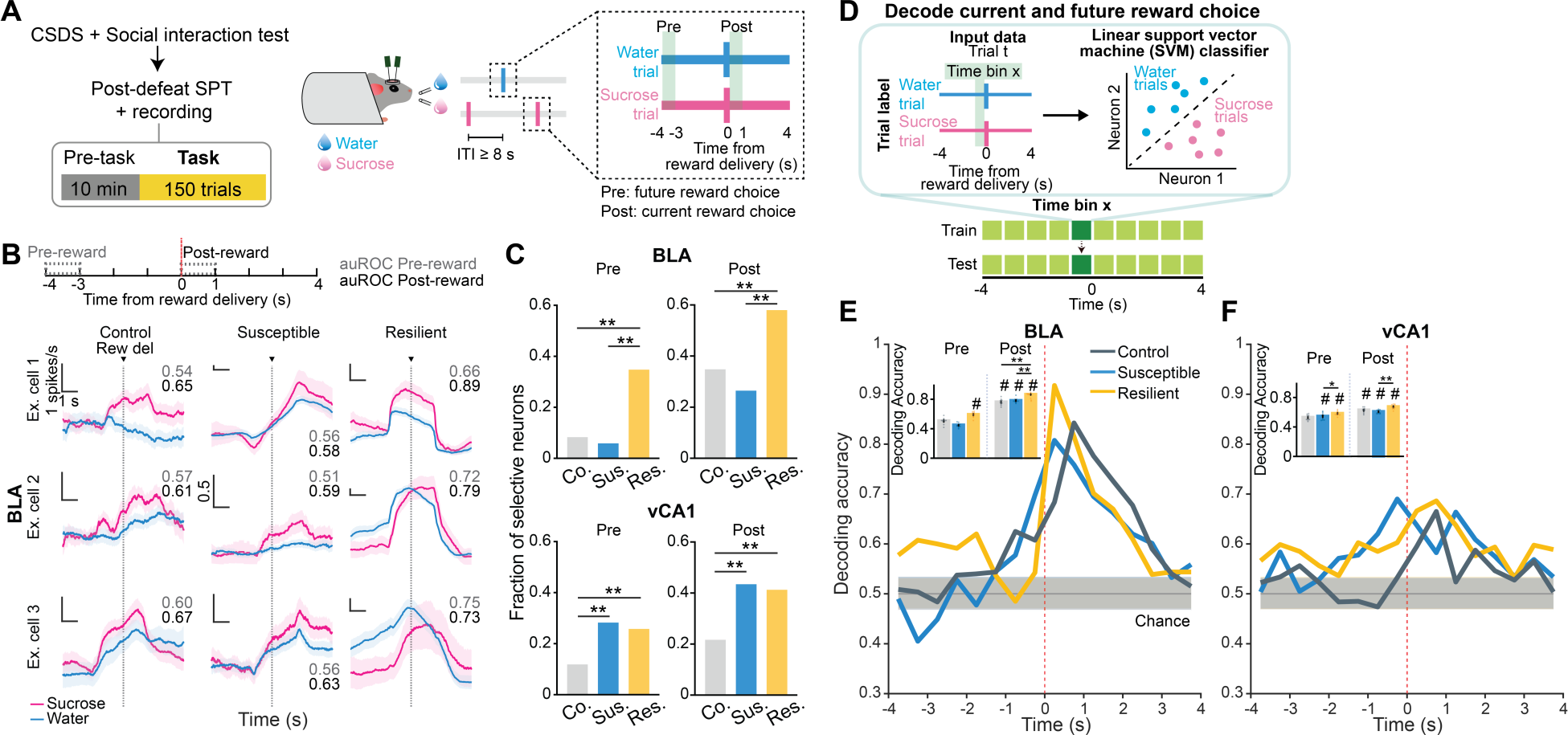
Enhanced representations of reward choice in stress resilient mice. (**A**) Schematic of SPT. **(B)** Trial-averaged firing rates in sucrose and water trials from example BLA cells, with respective auROC during Pre-reward (grey, -4 to -3s) and Post-reward (black, 0 to 1s). Scale bars: 1 spikes/ 1s unless otherwise stated. **(C)** In BLA, resilient group showed the highest fraction of selective neurons during both Pre-(Fisher’s exact tests, *P* < 0.0001) and Post-reward (Fisher’s exact tests, *P* < 0.001). In vCA1, both susceptible and resilient mice showed higher fraction of selective neurons than controls during Pre-(Fisher’s exact tests, *P* < 0.001) and Post-reward (Fisher’s exact tests, *P* < 0.0001). **(D)** Schematic of population decoding of current and future reward choices. Linear support vector machine (SVM) classifier was trained to distinguish between water and sucrose trials. **(E)** In BLA, resilient mice showed higher decoding accuracy than chance during Pre-reward, and the highest decoding accuracy among all groups during Post-reward (Kruskal-Wallis, *P* < 0.0001). **(F)** In vCA1, resilient mice showed higher decoding accuracy than susceptible mice during Pre-reward (Mann-Whitney, *P* = 0.045) and Post-reward (Kruskal-Wallis, *P* = 0.0011). Data are mean ± STD. # Significantly different from chance; * *P* < 0.05; ** *P* < 0.01.

### Intention-specific states in BLA as a unique susceptibility signature

We next examined the origins of anhedonic behavior in susceptible mice by analyzing the sequence of reward choices. We found that choices were not independent from each other, as the sequence could be described using a Markov model in which the probability of choosing water or sucrose depends on the choice in the previous trial. The Markov models of control and resilient mice were similar, both exhibiting a higher propensity to switch from water to sucrose and tended to repeat a sucrose choice more often than susceptible mice (Fig. 3A-B, Fig. S2C-F). In contrast, susceptible mice tended to switch away from sucrose rewards and stay on consecutive water choices. Given this, we asked whether there were unique neural signatures of the intention to switch or stay when the four possible sequences of consecutive reward choices (water-water; water-sucrose; sucrose-water; sucrose-sucrose) were considered.

**Fig. 3.**
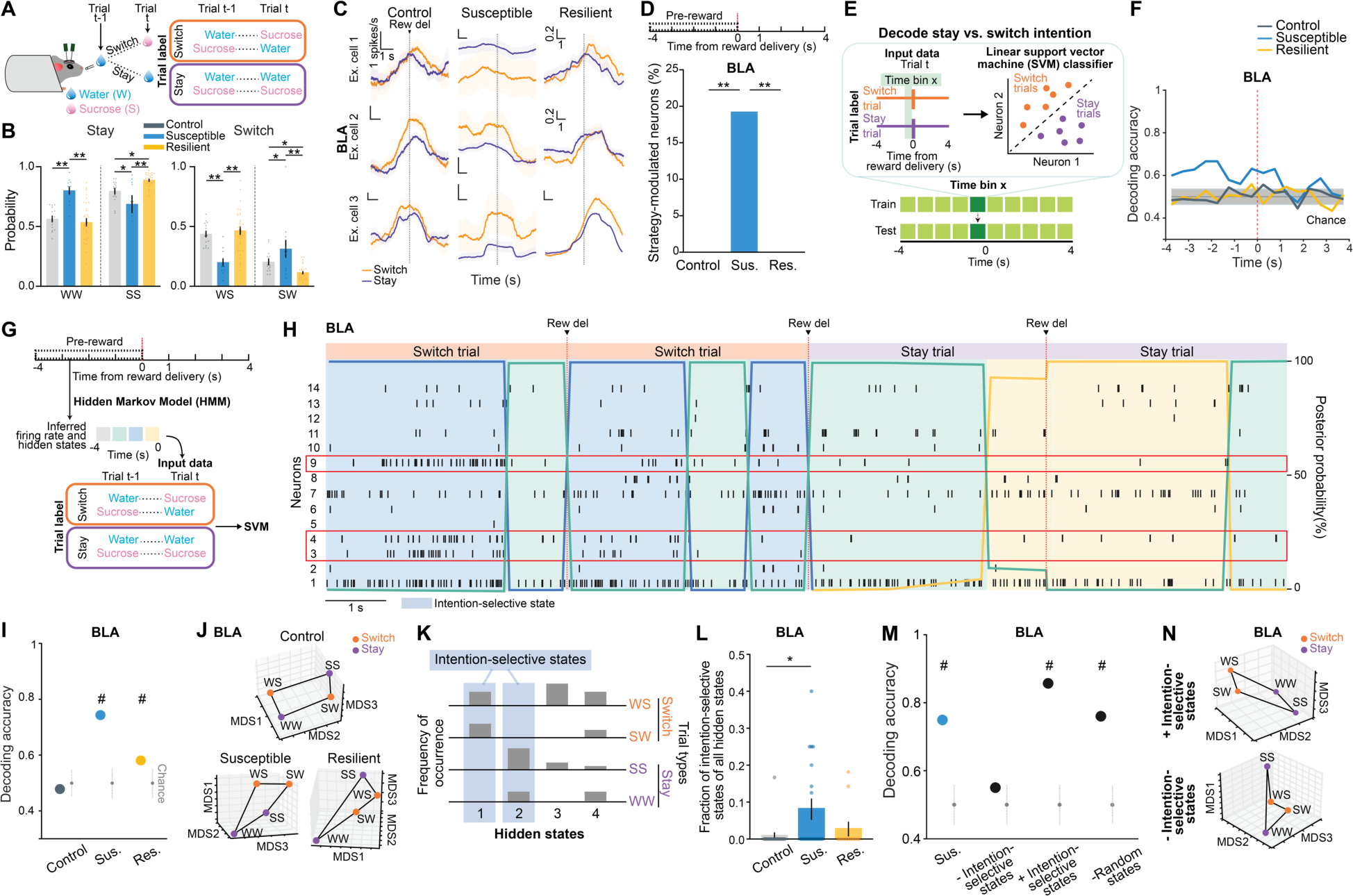
Intention-specific states in BLA as a unique susceptibility signature. **(A)** Schematic of switch/stay. **(B)** Susceptible mice had more water-water and sucrose-water trials (ANOVA, sucrose-sucrose, group: *F*_2,57_ = 14.84, *P* < 0.0001; water-water, group: *F*_2,57_ = 11.71, *P* < 0.0001; water-sucrose, group: *F*_2,57_ = 14.84, *P* < 0.0001; sucrose-water, group: *F*_2,57_ = 11.71, *P* < 0.0001). **(C)** Switch/stay trial-averaged firing rates from example cells. Scale bars: 1 spikes/ 1s unless otherwise stated. **(D)** Susceptible mice had more intention-selective neurons (Fisher’s exact test, *P* < 0.0001). **(E)** Schematic of switch/stay population decoding. **(F)** Susceptible mice showed greater-than-chance decoding accuracy. **(G)** Schematic of HMM. **(H)** Example spike rasters and hidden states. Neurons in red showed preferential firing during switch/stay trials. **(I)** Switch/stay could be decoded using HMM inferred firing rates in susceptible mice. **(J)** Switch/stay representations can be linearly separated in susceptible mice. **(K)** States that only occur in switch/stay trials are defined as intention-selective. **(L)** Susceptible mice had more intention-selective states (Kruskal-Wallis, *P* = 0.045). **(M)** Bidirectional modulation of trials containing intention-selective states bidirectionally changed decoding accuracy in susceptible mice. **(N)** Switch/stay representations before and after removal of intention-selective states. Data are mean ± SEM. # Significantly different from chance; * *P* < 0.05; ** *P* < 0.01.

Single cell analysis revealed that only in BLA of susceptible mice were there neurons that were differentially modulated based on the intention to switch or stay on the same reward as the previous trial (Fig. 3C-D). In addition, a population decoder could successfully distinguish between stay vs. switch trials in the seconds before reward was delivered in BLA of susceptible mice but not in the other groups (Fig. 3E-F) This was specific to BLA, as we did not observe striking switch vs. stay signatures at the population level in vCA1 (Fig. S2G).

This led us to hypothesize that specific population activity patterns existed in the BLA of susceptible mice in the seconds preceding a switch or stay decision. We identified population hidden states in the 4s Pre-reward period using Hidden Markov Models (*52–54*) (HMM, Fig. 3G-H, Fig. S2H) and confirmed that a linear decoder trained using inferred firing rates in HMM hidden states could most strongly distinguish between stay vs. switch trials in BLA of susceptible mice (and both vCA1 CSDS groups) (Fig. S2I-J), and the population representations of stay vs. switch were linearly separable in BLA of susceptible mice (Fig. 3I-J). Next, we identified hidden states that uniquely existed only in trials where mice either intended to stay or switch, which we termed intention-selective states (Fig. 3K-L, Fig. S2K, see Methods). We found that BLA of susceptible mice had a significantly higher fraction of these intention-selective states during the 4s Pre-reward period than controls (Fig. 3L). Removing trials that contained these states reduced decoding accuracy of stay vs. switch trials to chance levels (Fig. 3M) and altered the geometry of population representations of switch vs. stay trials (Fig. 3N). Considering only trials with intention-selective states improved the switch vs. stay decoding accuracy, while removal of random states did not impact decoding accuracy (Fig. 3M-N). Altogether, our results indicate that BLA neurons in susceptible mice evaluate future decisions with respect to their past choices (by representing switch/stay states), which may contribute to behavioral strategies that result in reduced number of sucrose rewards.

### Distinct patterns of spontaneous activity in the BLA of susceptible mice

We next asked whether distinct patterns of population activity could be detected in the BLA of susceptible or resilient mice, even without any overt stimuli or task demands. Altered resting state functional connectivity patterns in the AMY and HPC have been observed in depressed individuals, but the underlying neural dynamics remain unknown (*55, 56*). Mice were head-fixed, mimicking a mildly stressful experience in human imaging studies, but no other stimuli were provided. We identified general features shared among mice of the same group (control, susceptible, or resilient) by analyzing the geometry of neural representations (*57, 58*). The lack of behavioral labels makes it difficult to align the representations of different animals to compare their geometry. Hence we focused on the embedding dimensionality, which can be estimated without alignment (*57, 58*) (see Methods). This revealed a trend towards higher dimensionality in the BLA of susceptible mice compared to controls (Fig. S3A-E), suggesting a larger number of states visited, with each state spanning a different dimension. Quantification of states using HMM (Fig. 4A-C, Fig. S3F-L) demonstrated that in BLA, but not vCA1, susceptible mice showed a more distinct spatial structure of states than controls. Indeed, agglomerative clustering of these states revealed a significantly larger number of clusters in susceptible mice compared to control and resilient mice, suggesting that susceptible mice explored a greater number of distinct neural states. Consistent with this, average correlated BLA population activity across time was lower, and thus more variable, in susceptible mice (Fig. S3M-N). The greater number of distinct states in susceptible mice could not be attributed to an increased firing rate, which was lower in the BLA of susceptible mice compared to controls (Fig. S4O). Furthermore, across all mice, the number of distinct states was significantly correlated with behaviors used to assess anhedonia (Fig. 4D), suggesting a relationship between population hidden states in the BLA and classic readouts of anhedonia-related behavior.

**Fig. 4.**
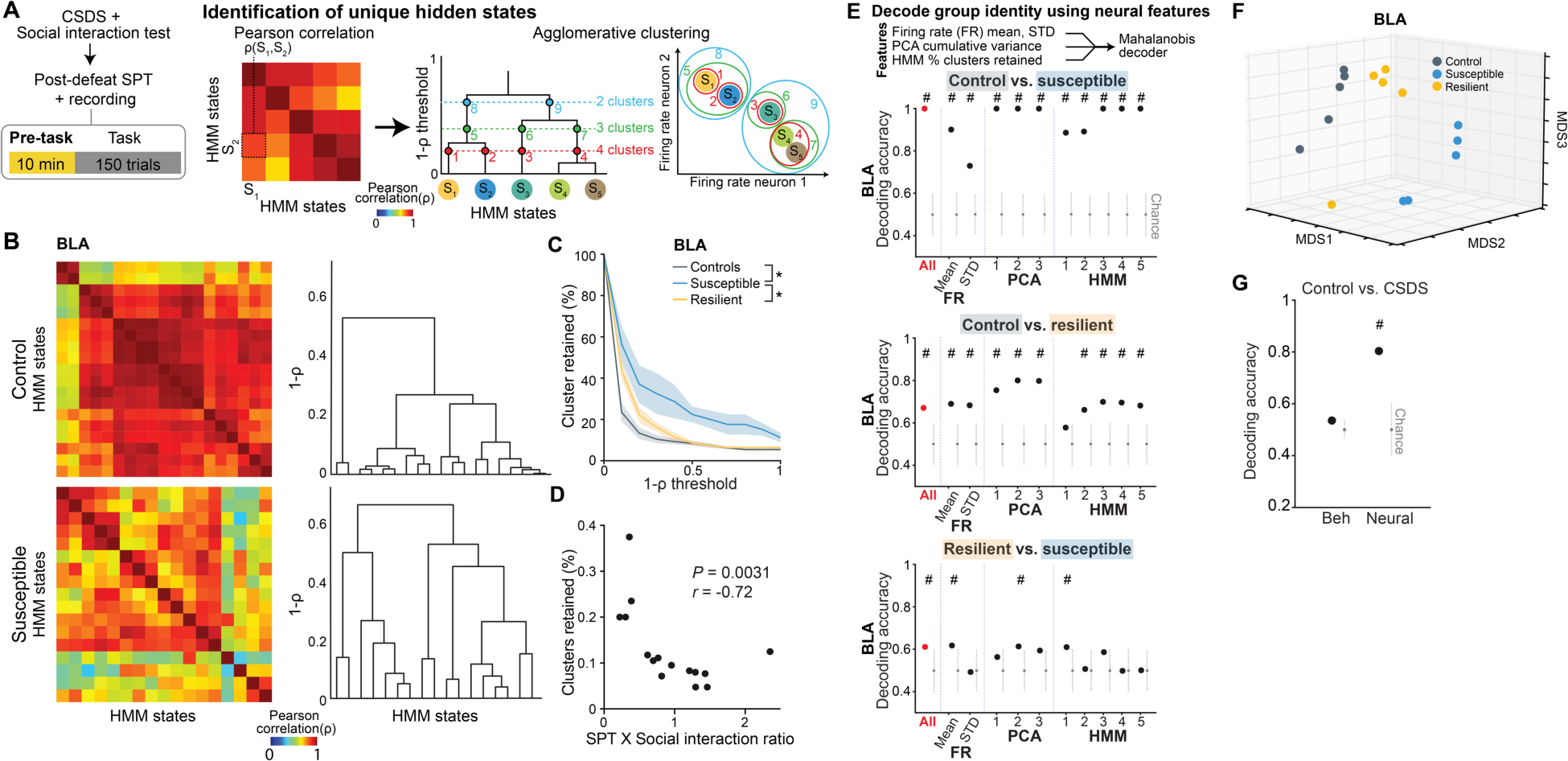
Distinct neural signatures of CSDS mice in the absence of task. (**A**) Schematic of analysis in pre-task. HMM was used to identify hidden states (S) and states similarity was assessed using agglomerative clustering. (**B**) Examples of states correlation heatmap from BLA of a control and a susceptible mouse and respective agglomerative clustering (dendrograms on right). (**C**) Susceptible mice had more distant hidden states in BLA (Mann-Whitney, control vs. susceptible *P* < 0.05 for all thresholds except 0; resilient vs. susceptible *P* < 0.05 for all thresholds except 0.4-0.6). (**D**) Proportion of clusters (1-π threshold = 0.5) was correlated with animals’ behavior (Spearman’s correlation). (**E**) Mahalanobis decoder trained on all neural features (“All”) could decode group identity better than chance in BLA. Feature importance in decoding was examined by systematic removal of each feature (subsequent columns). (**F**) MDS of neural features in BLA showed that controls were most distinct from susceptible mice. (**G**) Mahalanobis decoder trained on neural features was better at distinguishing control vs. CSDS mice than behavioral features. Data are mean ± SEM. Chance distributions are 2xSTD around theoretical chance level. # Significantly different from chance; * *P* < 0.05.

We next tested whether we could decode the group identity of individual animals, by training a classifier using neural features including firing rates (mean and standard deviation), PCA cumulative variance, and the fraction of agglomerated clusters across clustering thresholds. While each of these features alone could distinguish between control vs. susceptible mice to some extent (Fig. S3P), using all the feature sets in BLA, but not vCA1, allowed us to decode between all pairs of group identities with the highest accuracy (Fig. 4E, Fig. S3Q-R), with decoding of susceptible vs. control mice at 100% accuracy. When we visualized the geometry of the representations in individual mice, we found the greatest distance between control and susceptible mice in the BLA (Fig. 4F). Applying this analysis to the task period could also differentiate between control and susceptible mice (Fig. S4). Finally, we found that a decoder using these neural features during the stimulus-free pre-task period in BLA could better predict whether an animal was exposed to stress than a decoder using only behavioral features (Fig. 4G). This suggests that neural activity features in the BLA in the absence of any stimuli or task demands may be a more powerful biomarker for identifying a history of chronic stress than classic behavioral indices such as social avoidance and anhedonia-related behaviors.

### Rescue of dysfunctional vCA1-BLA activity and anhedonia by circuit-specific manipulations

We next explored neuronal circuit mechanisms that could contribute to these population-level signatures of resilience and susceptibility in the BLA. Specifically, we asked if we could rescue behavioral and physiological phenotypes of susceptibility via manipulation of inputs to BLA. We targeted vCA1 as 1) it provides dense input to BLA, 2) CSDS produces changes in representations of reward choice and intended task strategies in vCA1 of all mice, which may reflect a general adaptation to stress (see Fig. 2E-F, Fig. S2I), and 3) resilience was positively correlated with the strength of communication between vCA1-BLA for sucrose vs. water choices during the Pre-reward period (Fig. 5A-B). To test whether manipulation of vCA1-BLA inputs may modulate signatures of susceptibility in the BLA to influence anhedonic behavior, we increased the excitability of vCA1-BLA neurons using the chemogenetic actuator hM3Dq (*59, 60*) (Fig. 5C-D). We subjected mice to CSDS and recorded BLA and vCA1 activity and behavior in susceptible mice before and after injection of the hM3Dq activator clozapine-n-oxide (CNO). vCA1-BLA activation increased vCA1 firing rates (Fig. S5A-B, K) and modified population activity patterns in vCA1 (Fig. S5C-J). During the sucrose preference task, this manipulation enhanced vCA1-BLA correlations for sucrose vs. water choices during the Pre-reward period (Fig. 5E). In addition, we found that vCA1-BLA activation increased the decoding accuracy of reward choice Post-reward in both BLA and vCA1(Fig. 5F-G), a signature of enhanced reward choice representation that was associated with resilience (see Fig. 2E-F).

**Fig. 5.**
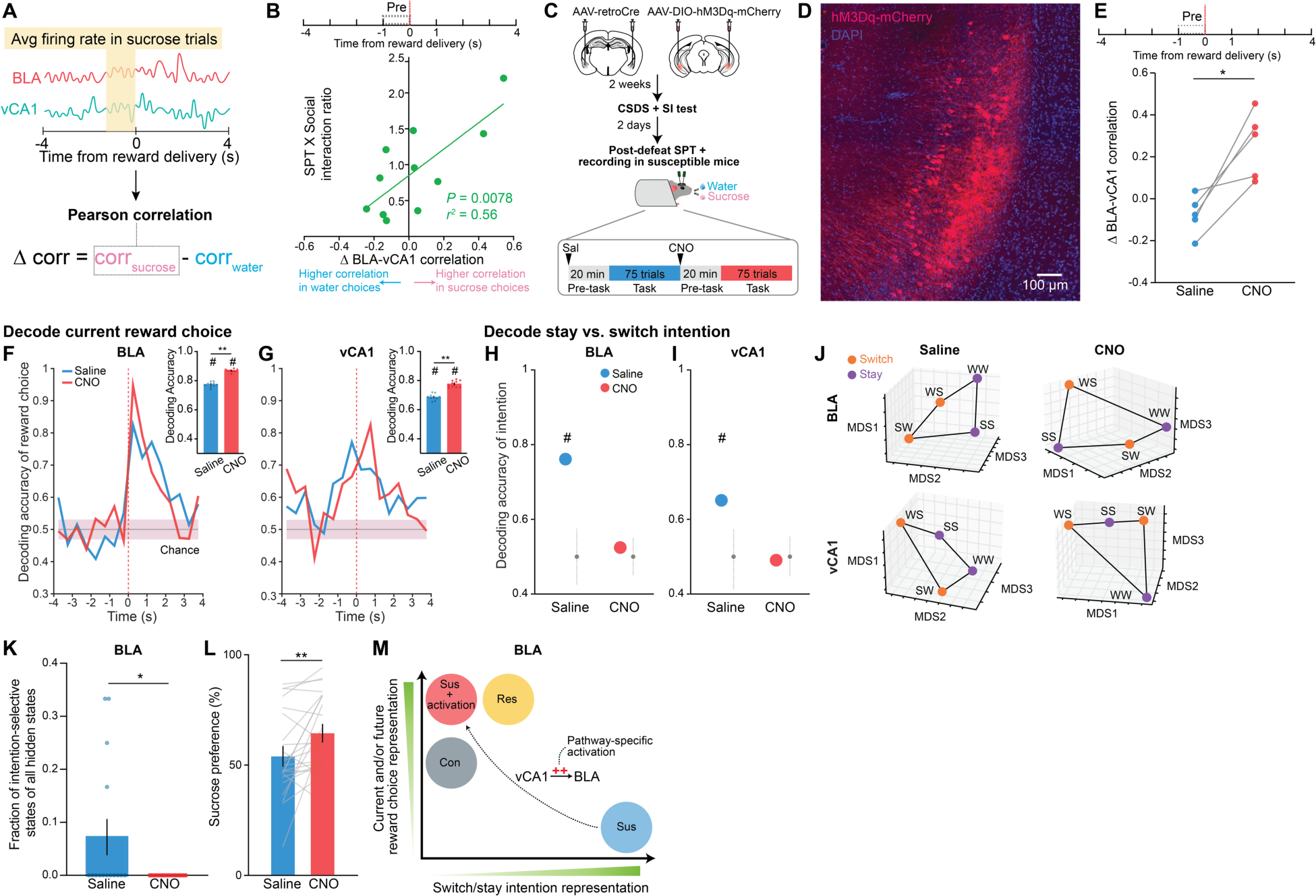
Rescue of dysfunctional vCA1-BLA activity and signatures of anhedonia by circuit-specific manipulations. (**A**) Schematic for analyzing vCA1-BLA interactions. (**B**) 1′BLA-vCA1 correlation Pre-reward was significantly correlated with animals’ behavior (Pearson correlation). (**C**) Schematic for chemogenetic activation of BLA-projecting vCA1 neurons. (**D**) Representative image of BLA-projecting vCA1 neurons transfected with hM3Dq-mCherry. (**E**) 1′BLA-vCA1 correlation was enhanced in CNO vs. saline (paired *t*-test, *t*_4_ = 4.22). (**F** and **G**) CNO increased decoding accuracy of current reward choice compared to saline (Mann-Whitney, *P* < 0.0001) in (**F**) BLA and (**G**) vCA1 (Mann-Whitney, *P* < 0.0001). Data are mean ± STD. (**H** and **I**) CNO reduced decoding accuracy of switch/stay trials to chance level in (**H**) BLA and (**I**) vCA1. (**J**) CNO altered the geometry of switch/stay representations such that they can no longer be easily distinguished. (**K**) CNO reduced the fraction of intention-selective states in BLA (Mann-Whitney, *P* = 0.043). (**L**) CNO increased sucrose preference (paired *t*-test, *t*_20_ = 2.39). (**M**) Summary schematic on the main findings. Data are mean ± SEM unless otherwise stated. # Significantly different from chance; * *P* < 0.05; ** *P* < 0.01.

We next asked whether manipulation of the vCA1-BLA pathway may reduce the occurrence of the unique intention-specific states we observed in the BLA of susceptible mice. Replicating our previous results, we found that during the saline period, we could decode stay vs. switch trials in these susceptible mice (Fig. 5H-J, Fig. S5L-M), however, activation of vCA1-BLA brought decoding accuracies to chance levels, changed the geometry of representations in the BLA such that switch and stay trials could no longer be linearly separated (Fig 5J), and decreased the fraction of intention-specific states (Fig. 5K). In other words, activation of vCA1-BLA pathway was able to reverse this population-level susceptibility signature in BLA.

Finally, we found that vCA1-BLA activation rescued behavioral indices of anhedonia, as it increased sucrose preference (Fig. 5L), increased lick rate discrimination index (Fig. S5N), and enhanced the proportion of sucrose stay trials (Fig. S5O-Q). No behavioral differences were observed between saline and CNO periods in mice infused with the control mCherry virus (Fig. S5R). In summary, these results show that activating vCA1-BLA pathway rescued both aberrant population dynamics in the BLA of susceptible mice and associated behavioral signatures of anhedonia (Fig. 5M).

## Discussion

Our study uncovers distinct stress resilience and susceptibility signatures in BLA population activity. Using Neuropixels recordings during both task-free and free-reward-choice periods and leveraging complementary analytic approaches, we identified novel population dynamics that underlie distinct features of the stress-induced anhedonic state. Furthermore, we successfully reversed behavioral and neural signatures of anhedonia through targeted modulation of the vCA1-BLA circuit.

By analyzing population dynamics during the sucrose preference test, we discovered a resilience signature characterized by heightened reward choice representations in the BLA before and during reward consumption. This enhanced reward choice perception or sensitivity may play a crucial role in reinforcing the behavioral choice that leads to the more rewarding option (i.e., choosing the sucrose reward) (*61, 62*), perhaps as a mechanism for adapting and coping with the experience of CSDS that ultimately results in a robust behavioral preference for sucrose. In contrast to resilient mice, susceptible mice exhibited reduced representations of current and future reward choices in the BLA, which may lead to under-estimation of the choice associated with the more rewarding option (*62*). Moreover, in the BLA of susceptible mice, we observed unique representations that reflect the intention to switch or stay on the previously chosen reward. This heightened evaluation of future choices with respect to the past in the BLA of susceptible mice is reminiscent of rumination-like states commonly observed in individuals with depression, such as repetitive thinking of past choices and/or future decisions (*63, 64*).

By analyzing the neural activity patterns in the absence of any task or stimuli, we found an enhanced exploration of distinct neural states in the BLA of susceptible mice. This may reflect the emergence of intrusive activity patterns in the BLA, like intrusive thought patterns observed in depressed patients (*65, 66*). Notably, we found that features of neural activity in the BLA during a task-free period were more effective than behavioral readouts in distinguishing between control mice and those with a history of CSDS. This suggests the intriguing possibility that resting-state activity patterns in the BLA hold significant potential as a powerful biomarker for predicting individuals who have experienced a stressful event.

Finally, susceptible mice showed reduced vCA1-BLA correlations during higher value sucrose reward choice trials, potentially driving anhedonic behavior. Consequently, chemogenetic activation of BLA-projecting vCA1 neurons in susceptible mice increased inter-regional communication between BLA and vCA1, increased representations of current reward choice in both BLA and vCA1, reduced the rumination-like over-representation of the intention to stay or switch in the BLA, and reduced anhedonia-related behavior. Our data suggests that vCA1 may encode an adaptive coping response to stress, as reward choice representations were enhanced in both resilient and susceptible mice in comparison to controls. This information from vCA1 may be funneled to the BLA via direct vCA1-BLA projection to help further distinguish between resilient and susceptible mice in the BLA. Specifically, in resilient mice, as BLA is more effectively interacting with vCA1, it may be incorporating this adaptive stress response from vCA1 to generate signatures of resilience, such as enhanced reward choice representations. In contrast, in susceptible mice, this vCA1-BLA interaction is weaker, thus, BLA may not be efficiently using this adaptive information from vCA1, resulting in neural activity patterns that promote susceptibility, such as reduced representations of reward choices and emergence of intention-selective states.

While dysfunction in dopaminergic systems has been implicated in motivational changes in depression and chronic stress (*3, 67–71*), this work provides crucial evidence for a role for the vCA1-BLA circuit in modulating stress-induced behavioral phenotypes. By demonstrating that targeting the vCA1-BLA circuit could normalize neural dynamics associated with susceptibility and promote those associated with resilience in the BLA, our findings shed light on how dysfunction in this circuit may contribute to stress-induced maladaptive states. Moreover, these results highlight the vCA1-BLA circuit as a promising target for neuromodulation in mood disorder treatments and open new avenues for potential therapies to address stress-induced pathologies more effectively.

## Acknowledgements

We thank L. Frank, V. Namboodiri, V. Sohal, J. Biane, A. Klein, and J. Bratsch-Prince for comments and discussion.

## Funding

Canadian Institutes of Health Research Postdoctoral Scholarship (FX)

Brain and Behavior Research Foundation Young Investigator Award (FX)

Ray and Dagmar Dolby Family Fund (FX, MAK)

Simons Foundation (VF, SF) Neuronex NSF1707398 (VF, SF)

Gatsby Charitable Foundation GAT3708 (VF, SF)

Swartz Foundation (VF, SF)

National Institute of Mental Health R01 MH108623, R01 MH111754, R01 MH117961, R01 MH125515 (MAK)

National Institute on Deafness and Other Communication Disorders R01 DC019813 (MAK)

One Mind Rising Star Award (MAK)

Human Frontier Science Program RGY0072/2019 (MAK)

Esther A. and Joseph Klingenstein Fund (MAK)

Pew Charitable Trusts (MAK)

McKnight Memory and Cognitive Disorders Award (MAK)

## Author contributions

Conceptualization: FX, MAK

Methodology: FX, VF, SF, MAK

Investigation: FX, VF Visualization: FX, VF

Funding acquisition: FX, VF, SF, MAK Writing – original draft: FX, MAK

Writing – review & editing: FX, VF, SF, MAK

## Supplementary Materials

## Materials and Methods

### Mice

All procedures were conducted in accordance with the NIH Guide for the Care and Use of Laboratory Animals and institutional guidelines. Adult (8-12 weeks old) male and female C57BL/6J mice were supplied by Jackson Laboratory. Adult (5-6 months old) CD1 retired male breeder mice were supplied by Charles River. All mice were kept on 12-hour light/dark cycle, and all experiments were conducted during the light phase. We performed recordings in 60 mice for the original dataset, including 45 CSDS mice (30 males, 15 females) and 15 control mice (10 males, 5 females). A separate cohort of 41 CSDS mice (males) underwent chemogenetic manipulation experiments. Twenty-three of the mice received AAV-DIO-hM3Dq virual micro-infusion and 18 mice received AAV-DIO-mCherry infusion. From the hM3Dq group, we performed recordings in 7 of the susceptible mice.

### Surgery

#### Head bar and craniotomy surgery

One week prior to lick training, head bar surgeries were conducted on all mice (8-9 weeks old). Similar to previously described protocol (*46*), mice were anesthetized with 1.5% isoflurane with O_2_ flow rate of 1 L/min, and head-fixed in a stereotaxic frame. A custom-made titanium head bar was then attached to the skull using Metabond adhesive cement (Parkell, Edgewood, NY). Possible recording sites (see Neuropixels section) were stereotaxically marked using a permanent marker on the skull surface, and skull was covered using silicon (Smooth-On, Macungie, PA). Three days prior to Neuropixels recording, craniotomy surgery was performed, where under anesthesia, craniotomies were made at previously marked coordinates. Skull surface was covered with Kwik-Sil (World Precision Instruments, Sarasota, FL).

#### Viral micro-infusion surgery

For mice that underwent chemogenetic manipulations, adult males (8-9 weeks old) received viral micro-infusion in the same surgery as head bar attachment, similar to previously described protocol (*46*). Specifically, AAV8-hSyn-DIO-hM3D(Gq)-mCherry (Addgene, 44361-AAV8, 2.9 x 10^13^ vg/ml) or AAV8-hSyn-DIO-mCherry (Addgene, 50459-AAV8, 1.0 x 10^13^ vg/ml) was micro-infused in vCA1 bilaterally (500 nL per hemisphere, -3.52 mm AP, ±3.1 ML, -4.2 [150 nL], -4.1 [200 nL] and -4.0 [150 nL] DV, from bregma according to Paxinos and Franklin [2011]), and AAV2retro-CAG-Cre (UNC Vector Core, Ed Boyden’s stock, 4.1x10^12^ vg/ml) was micro-infused in BLA bilaterally (500 nL per hemisphere, -1.80 mm AP, ±3.1 ML, -5.0 [150 nL], -4.8 [200 nL] and -4.6 [150 nL] DV). Viral vectors were delivered using Nanoject 3 (Drummond Scientific, Broomall, PA). The needle was held in place for >5 min after infusion at each DV, and for 10 min after the last DV. Following viral micro-infusion, head bar was attached to the skull as described above.

### Behavior

#### Chronic social defeat stress (CSDS)

CSDS was conducted similar to previously established protocol (*6*). Briefly, CD1 male mice were singly housed upon arrival for >1 week and were then pre-screened for aggression over 3 consecutive days. Each day, a CD1 mouse was placed in a cage with a new screener BL/6 mouse for 3 min. An aggressive CD1 mouse is defined as one that attacked the BL/6 mouse within the first min over a minimum of 2 consecutive days. Only aggressive CD1 mice were used in defeats and social interaction tests. Defeats occurred over 10 days, where each day, a BL/6 mouse was introduced to a novel CD1 mouse’s cage for 10 min. Defeats were terminated early if severe injuries on BL/6 mice were observed. After 10 min, a clear plastic divider with perforations was placed in the middle of the defeat cage for 24 hours, to physically separate the BL/6 and CD1 mice while allowing visual and odor cues to transmit and reinforce the defeat experience during co-housing. After the 10^th^ day defeat, BL/6 mice were singly housed in new cages (without CD1s) for 24 hours prior to social interaction test. For female defeats, female BL/6 mice were first coated with urine from other aggressive CD1 male mice (not used in defeats) prior to being introduced to the defeat CD1 mouse cage (*72*), in order to minimize mounting behavior and maximize defeats. Female defeats were terminated early if mounting was observed.

#### Social interaction test

Social interaction test took place 1 day after termination of CSDS. BL/6 mice were habituated to the social interaction test room for 1 hour prior to test. The test was performed under red light (10 lux) in a test arena (custom made, 42 cm (w) × 42 cm (d)× 42 cm (h)) in a sound attenuation chamber. During the first phase of the test, the BL/6 mouse was introduced to the test arena with an empty enclosure (10 cm (w) × 6.5 cm (d) × 42 cm (h) at one end for 2.5 min, and its activity patterns were tracked using Ethovision (Noldus Information Technology, Leesburg, VA). At the end of 2.5 min, the mouse was placed back in its home cage, and the empty enclosure was replaced with a second enclosure containing a novel aggressive CD1 that had not been used in defeats. The BL/6 mouse was put back in the test arena for another 2.5 min. Social interaction ratio, as a measure for social avoidance, was calculated as the time spent in the interaction zone (14 cm × 24 cm) with the aggressor present vs. absent. The lower the social interaction ratio, the more socially avoidant the animal was.

#### Head-fixed sucrose preference test

Following recovery from head bar surgery, mice were habituated to the experimenter and the head-fixed setup for 15 min a day for a week. After habituation, mice were water-restricted to ∼85-90% their ad lib body weight and were trained for 3 days to lick on the custom designed dual-spout head-fixed reward delivery apparatus. On day 1, mice were introduced to 1 lick spout, where sucrose rewards (10% sucrose, ∼3.5 ml each) were intermittently delivered upon licking (i.e., reward were lick-contingent) with 8 seconds inter-trial interval (ITI), with maximum 150 rewards per session. Sucrose rewards were delivered using a solenoid-gated gravity feed. Licks were detected using a piezo element (SparkFun, Boulder, CO). Stimulus delivery and sensor reading were controlled using a custom Arduino MEGA board and recorded using CoolTerm software. On day 2 and 3, mice were introduced to 2 lick spouts, one on each side of the mouse, separated by ∼50 degrees. Sucrose rewards were delivered in both spouts upon licking with 8 seconds ITI. The goal was to teach mice that rewards were delivered from both spouts. Thus, if a mouse showed preference for spout on one side, that spout was temporarily removed so the mouse can learn to lick from the other spout. Once the animal showed similar preference for both spouts, lick training was completed and pre-defeat sucrose preference test (SPT) was initiated on the following day. SPT occurred over the course of 2 consecutive days, where one spout delivered water and the other delivered sucrose. Rewards were delivered upon licking with 8 seconds ITI and maximum 150 rewards in total per day. The spout designation was randomized across mice on day 1 and counterbalanced on day 2. Sucrose preference was calculated as the averaged % sucrose rewards obtained across 2 days. Upon completion of day 2 of pre-defeat SPT, mice were taken off water restriction and housed in social defeat room for 3 days before CSDS began. Post-defeat SPT was performed using the same protocol, with the addition of Neuropixels recording. Sucrose preference post-defeat was used for all behavioral analysis.

### Neuropixels recording and data pre-processing

#### Recording

Mice were head-fixed to the SPT apparatus without lick spouts present. Kwik-Sil was removed from skull surface. Prior to insertion, Neuropixels 1.0 probes (IMEC, Heverlee, Belgium) were first coated with DiI, DiO, or DiD dyes (ThermoFisher Scientific, Waltham, MA) and allowed to dry. Probes were inserted at ∼1 mm/min to the target coordinate using Sensapex manipulators (Oulu, Finland). Possible probe targets and their coordinates are as follows: amygdala (-1.71 mm AP, -0.28 mm ML, -6.5 mm DV, at 31.3 degrees ML), ventral hippocampus (-3.9 mm AP, -2 mm ML, -4.5 mm DV, at 25.8 degrees ML), frontal cortex (+1.77 mm AP, 0 mm ML, -5.7 mm DV, at 9 degrees ML; +2.60 mm AP, -0.5 mm ML, -4.5 mm DV, at 9 degrees ML), and midline thalamic and hypothalamic regions (-3.80 mm AP, 0 mm ML, -5 mm DV, at 7.6 degrees ML; -1.90 mm AP, -0.15 mm ML, -4.3 mm DV, at 4.4 degrees ML). One or two probes were inserted per session per mouse. Simultaneously recorded probes were coated in the same color of dye but spaced at least several hundred microns apart to allow for unambiguous identification. Different colors of dyes were used across days to help differentiate probe tracks. After a probe reached targeted DV, it was left in place for 10 min prior to the start of recording and SPT. Neuropixels action potential signals were recorded using Neuropixels acquisition system and SpikeGLX software, at 30,000 Hz with gain = 500. Behavioral signals were recorded using a separate data acquisition board (National Instruments, Austin TX), along with a synchronization signal that was also recorded by Neuropixels to help synchronize clocks between different data streams. After each session of SPT, probes were slowly removed from the brain and skull was covered with Kwik-Sil. Probes were cleaned using Tergazyme solution (1%, Alconox) overnight and rinsed using deionized water before reusing or storage.

#### Histology and probe track registration

At the end of experiments, mice were transcardially perfused with 1x PBS followed by 4% paraformaldehyde solution. Brains were fixed overnight at 47C, and then transferred to 30% sucrose solution for 48 hours. Brains were sectioned coronally using a microtome (Leica SM2000) at 50 um thickness and mounted on glass slides with Fluoromount G with DAPI (Southern Biotech, Birmingham, AL). Images were obtained using a confocal microscope (Nikon Ti2-E Crest LFOV Spinning Disk/ C2 Confocal) with 20X objective. Probe tracks were traced using the AllenCCF toolbox (https://github.com/cortex-lab/allenCCF).

#### Spike-sorting

Neuropixels action potential signals were pre-processed and spike-sorted offline using Kilosort 2 (*73*), and after sorting, the clusters were manually validated using Phy (*74*). Only well-isolated clusters (putative single units that are classified as “Good” using Phy) were analyzed. All other clusters, including multi-unit activity and noise, were not analyzed.

### Data analysis

Animals were allowed to freely choose reward types after 8s ITI had passed between trials, by licking at the spout of their choice. Reward deliveries were lick-contingent. Trial types were defined as ± 4s time window around the time of reward delivery. For all analysis, only mice with at least 5 neurons in the region of interest were used.

### Behavioral data analysis

#### Behavioral classification of mice

The relationship between sucrose preference and social interaction ratio was assessed using Pearson’s correlation. To classify CSDS mice into subtypes, we applied unsupervised K-means clustering using both behavioral metrics, sucrose preference and social interaction ratio. The optimal number of clusters was determined by evaluating cluster numbers from 2 to 10 and maximizing the Silhouette score.

#### Lick analysis

Lick rasters were generated by binning licks using 0.02s bin size. Lick rates were calculated using 0.1s bin size and averaged across trials per mouse for each reward type. Lick rate discrimination index (DI) was calculated as the difference between lick rates for sucrose vs. water trials divided by the sum of the two.

To take into account reward history and assess how it affects current behavior, we further divided sucrose and water trials into sucrose/sucrose (SS), water/sucrose (WS), water/water (WW), and sucrose/water (SW) trials, (previous/current reward). The first trial of each session was discarded as it had no prior trial. In order to assess the probability of each trial type independent of the animal’s overall sucrose preference, we normalized the number of trials to the total number of previous trials of a specific type. For example, we defined the overall transition probability from a water trial to a sucrose trial as *P(*WS) = *P(WS)* / *(P(WW) + P(WS))*, and from a sucrose trial to a sucrose trial as *P(SS)* = P(SS) / *(P(SW) + P(SS))*, where *P(XY)* is the transition probability from reward X to reward Y. We normalized the transition probabilities such that *P(WW)+P(WS)* = 1, and *P(SS)+P(SW)*=1. Using this normalization, if *P(SW)* is not significantly different from *P(WW)*, this would suggest that the current water reward choice is independent of the previous reward, because of the probability of switching from sucrose or staying on water is the same; otherwise, the current reward choice is dependent on the previous reward (i.e., reward choices could be modeled as a first-order Markovian process).

We also computed the proportion of each of the 4 trial types normalized to the total number of trials per session, to assess how much each trial type contributes to the overall session. In this case, the % of trials of each of the 4 trial types were computed per session and averaged across sessions for each mouse. As the number of trials may be influenced by each animal’s innate preference for different rewards, we computed the chance probability of the occurrence of each trial type by calculating the joint probability of the previous and current trial. For example, *chance P(SW) = P(S) x P(W).* The number of trials was then subtracted by the chance level in each mouse (“Number of trials chance removed”). SS and WW trials were combined when analyzing stay trials, and SW and WS trials were combined when analyzing switch trials. The preference between stay and switch trials in each mouse was calculated as % stay trials - % switch trials. To quantify the number of consecutive trials, we first obtained the average number of consecutive trials per trial type (sucrose or water) per session and then averaged across sessions for each mouse.

#### Decoding group identity using behavioral features

To examine whether group identity could be decoded using behavioral data, we defined a Mahalanobis-like binary decoder. Specifically, for each mouse, we considered 4 behavioral features: lick rate DI during Pre-reward, Post-reward, sucrose preference, and SI score. Considering two groups at a time, we defined and constructed a Mahalanobis binary decoder to assign a single testing mouse to one of the two groups in the behavioral feature space. The input to the binary classifier consisted of an *N* x *F* training matrix and a 1 x *F* testing matrix, where N represented the total number of training mice between the two classes, and *F* = 4 represented the total number of features. In each cross-validation (CV), we first balanced the number of mice in each group by randomly subsampling the minimum number of mice between the groups. Next, we randomly selected one mouse as the testing sample and used the remaining mice as the training set, for a total of 1000 CVs. We defined a Mahalanobis-like distance in the feature space as the Euclidean distance between the testing mouse and the centroid of the training groups, divided by the variance along the distance direction. The testing sample was assigned to the group identity with the minimum Mahalanobis-like distance. The performance of the decoder was evaluated by calculating the fraction of correct classifications out of the total 1000 CVs, and the entire procedure was repeated for all possible pairs of the three groups (i.e., control, susceptible, and resilient mice).

### Single neuron analysis

#### Firing rate

For task period, spike trains were aligned at the time of reward delivery (time 0) and neurons within the same region were pooled across animals of the same group to construct pseudo-populations. Only neurons with at least 10 trials per trial type (sucrose and water) were included. For peri-stimulus time histograms, spikes were binned at 10ms resolution, z-scored to - Pre-reward (-1 to 0s), and smoothed with a 50ms moving average filter. For analysis of raw firing rates, spikes were binned at 500ms resolution.

#### Reward-modulated neurons

Analysis was performed using pseudo-population and only neurons with at least 10 trials per trial type (sucrose and water) were included. Reward-modulated neurons were identified based on Wilcoxon rank-sum test, comparing the distribution of firing rates during 1s Post-reward (0 to 1s) vs. 1s Pre-reward period (-4 to -3s) across trials. A cell is deemed reward-modulated if Post-reward epoch is significantly different from Pre-reward after false discovery rate (FDR) correction across all recorded neurons, with a significance threshold of *P* < 0.05. A Chi-squared test was used to perform statistical comparisons between fractions of reward-modulated neurons in CSDS vs. control mice for sucrose and water trials.

#### Intention-modulated neurons

Analysis was performed using pseudo-population and only neurons with at least 10 trials per trial type (switch and stay) were included. Intention-modulated neurons were identified using a similar method as reward-modulated neurons. Mice with less than 5 neurons in regions of interest were excluded. In this case, a cell is deemed intention-modulated if the distribution of firing rates during the 4s pre-reward period (-4 to 0s) in switch trials is significantly different from stay trials, as identified using Wilcoxon rank-sum test followed by FDR correction across all neurons in that group (*P* < 0.05). As the fraction of neurons was small and did not meet the criteria for using Chi-squared test, Fisher’s exact tests were used to perform statistical comparisons between % of intention-modulated neurons across groups.

#### auROC

Analysis was performed using pseudo-population and only neurons with at least 10 trials per trial type (sucrose and water) were included. Mice with less than 5 neurons in regions of interest were excluded. Reward-choice-selective cells were identified (*75, 76*), and the magnitude of the selectivity was quantified, using the area under the receiver operating characteristics (auROC) method, which compares single-neuron firing rates between trial types, across levels of response thresholds for each time bin. Spikes were binned at 500ms resolution. Shuffled distributions were computed for each time bin by randomly shuffling trial type 10 times per neuron. A neuron is deemed reward-choice-selective if its auROC is > 2 x STD of shuffled distribution for that neuron. The fraction of selective neuron in a region was calculated as # of selective neurons / total # of neurons. Differences in the fraction of selective neurons across groups were assessed using Fisher’s exact tests.

### Population analysis

#### Analysis of embedding dimensionality

Principal component analysis (PCA) was used to evaluate the embedding dimensionality of population activity of simultaneously recorded neurons over time. The method aims to identify how much variance of the population representation in the firing rate space is accounted for by each principal component. We chose this method because the pre-task period lacks behavioral labels. PCA has the advantage of allowing us to compare neural data between animals because the method is invariant for rotations and global stretching, transformations normally needed to align a neural representation of one subject into another. We examined the activity of each neuron in 1s bins during the 6 min time window (min 2-8) within the 10 min pre-task recording period, resulting in 360 bins. The ensemble activity across these bins can be represented as a geometrical object in the firing space, with each axis representing the firing rate of a neuron and each point representing the ensemble’s activity in a time bin. We calculated the embedding dimensionality of this geometrical object for each mouse. We only included mice with at least 5 simultaneously recorded neurons in the region of interest during the pre-task recording. We randomly selected 5 neurons for each mouse and calculated the z-scored firing rate matrix *N* x *T*, where *N* is the number of neurons, and *T* is the number of time bins. We applied PCA to this matrix and determined the cumulative curve of the variance explained by each principal component (PC). We repeated this procedure 1000 times and averaged the results across the subsamples for each mouse. Our goal was to compare cumulative variance curves across groups and determine if a group had a higher cumulative value at *M* PCs (*M* :: 5), indicating a lower dimensionality of the geometrical object. We subsequently used the cumulative variance values for the first 3 PCs as features to decode the group identity.

We also assessed the Participation Ratio (PR), which is a normalized measure of dimensionality based on the full distribution of PCA eigenvalues (i.e., how much variance is explained by each principal component), and it is defined as follows,

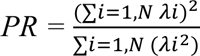

where *λ_i_* are the eigenvalues of the covariance matrix of the neural acidity, and *N* = 5. If only one eigenvalue explains all the variance (*λ_i_* ≠ 0 for *i* = 1 and *λ_i_* = 0 for all *i* ≥ 2), then *PR =* 1. On the other hand, if all eigenvalues are equal, the dimensionality is maximum, *PR* = *N* (*77, 78*).

During the task period, the same analysis was repeated during the 1s of Pre-reward and Post-reward periods, using z-scored firing rate with 0.2s bins (5 bins for each period).

#### Hidden Markov Model

We used hidden Markov models (HMMs) to identify patterns of population activity in the time series, where each pattern corresponds to a specific neural state that is not directly measurable (*52, 54, 79*). We fitted an HMM separately for each mouse for the pre-task and task period. For the DREADD dataset, HMMs were fitted for saline and CNO periods of each mouse separately. To perform model fitting, we employed the software framework developed by the Linderman Lab (https://github.com/lindermanlab/ssm). To prepare the data for the HMM analysis, we binned the 6 min pre-task recordings of each session into 1s bins, resulting in 360 bins. We computed the spike count of each neuron in each bin. The input data for the HMM consisted of an *N* x *T* matrix, where *N* represents the total number of simultaneously recorded neurons in the session, and *T* represents the total number of time bins.

For the analysis during task in the Pre-reward and Post-reward periods, we computed the spike count in 0.2s time bins. We fitted separate HMMs for the Pre-reward and Post-reward periods for sucrose and water trials. To accomplish this, we concatenated the M trials within a single session and arranged the input data in an *N* x *T* x *M* matrix, where *T* = 5. We chose bin size of 0.2sec, because this bin size balanced the inference of maximum possible transition states and total spike count used to fit HMMs.

For decoding of switch vs. stay using HMM states, we focused on the 4s pre-reward period. Spike counts were binned using 1s bin size, and concatenated across the 4s window of all trial types. This resulted in an *N* x *T* x *M* input matrix, where *T* = 4, and *M* represents the total number of recorded trials in the session. Consistent with previous analyses, we retained only sessions with at least 5 simultaneously recorded neurons.

Given the recorded (observed) spike count over time, we modeled the neuronal activity as a Poisson process, where the mean value depends on the current neural state. We represented the probability of observing the spike count vector n(t) of N neurons at time bin *t*, given the hidden neural state *S_t_* = *j*, as a multivariate Poisson process: *P (n_t_ | S_t_ = j) ∼ Poisson(Λ; n_t_)*. Here, *Λ* = {*λ_1_, λ_2_, … λ_N_*}, and *λ_i_* represents the estimated mean activity for the i^th^ neuron in state *j*. The vector *Λ* corresponds to the column of the *N* x *K* “emission matrix” *E*, which provides the firing rates or activation probabilities of observing a specific neuronal pattern when the population activity is in a particular state.

We assumed the dynamics of the neural states to evolve according to a first-order Markovian process, where the probability of transitioning from one state to another depends only on the current state. This process is summarized by the *K* x *K* “transition probability” matrix *T*. Additionally, we incorporated an initialization vector *A*, which provides the probability of starting in each state. The HMM was fully described by the set of parameters {*E, T, A*}, which were inferred by fitting the model to the recorded neuronal spike counts (*80*). We used the Baum-Welch expectation-maximization algorithm to update the model parameters and maximize the likelihood of the observed data. For each time series, we fitted 5 models with a maximum of 100 iterations for each value of the total number of states ranging from 2 up to 50, using randomized initial conditions. The model with the smallest Akaike Information Criterion (AIC) score was retained as the best model for further analyses (*52*). Subsequently, we used the Viterbi algorithm to estimate the most likely sequence of states over time.

#### Agglomerative clustering analysis

To better characterize the spatial structure of the hidden states, we examined the pairwise correlation between the inferred activity of the states. For state 1 with an activity vector *X* = (*x_1_, x_2_, …, x_N_*), where *x_i_* represents the activity of neuron *i*, and state 2 with an activity vector *Y* = (*y_1_, y_2_, …, y_N_*), we computed the Pearson correlation coefficient *ρ(X,Y)* to assess the distance between the states in the neuronal activity space. We calculated the correlation coefficients for all pairs of states and stored them in an *N* x *N* correlation matrix *K.* Subsequently, we performed agglomerative clustering on the correlation matrix.

Specifically, we defined a new distance matrix *D* as 1 - *K*, where 1 is an *N* x *N* matrix of ones. This matrix served as the input to the agglomerative clustering algorithm, which iteratively combines states to define new clusters according to the pairwise distance. The algorithm initialized each state as a separate cluster with minimum distance (maximum correlation) and iteratively merged two clusters *v* and *u* with the smallest distance into a new cluster. The new distance *d* assigned to the agglomerated clusters was defined as *d(u,v) = max(dist(u[p], v[q]))*, where *p* and *q* represent all the points in the merged clusters *u* and *v*, also known as Farthest Point Algorithm. Agglomerative clustering has the advantage of producing a hierarchical structure of clusters, which we represented as a dendrogram. This hierarchical representation allowed us to examine the relationships and similarities between states, specifically how neural states may be nested differently within large clusters in different groups. Importantly, agglomerative clustering does not require any assumption regarding the total number of clusters. It iteratively merges the closest states and clusters until all states are merged into one final cluster. We performed the clustering analysis separately for each mouse, visualizing the results with a dendrogram that summarizes the merging of clusters at different levels of distance, ranging from 0 (original states) to 1 (a single cluster).

After examining the clusters, we counted the total number of clusters at different levels of distance, or thresholds, where the higher the levels of distance, the lower the number of clusters, until reaching only one cluster at the highest distance. We assessed the curves of the number of total clusters and the proportion of total clusters retained relative to the total number of states as a function of thresholds. Comparing these curves between two groups, a higher number of states at the same threshold value indicates a greater degree of dissimilarity among the inferred states. We retained the proportion of total clusters along these curves from a threshold of 0.1 up to 0.5, resulting in a total of five features that were subsequently used in the decoding of group identity. We applied the clustering analysis to the pre-task activity using the previously inferred states described in the “Hidden Markov Model” section, as well as to the Pre-reward and Post-reward task periods for water and sucrose trials separately.

#### Correlation of population activity across time

To examine how variable population activity was across time during the pre-task period, we performed Pearson’s correlation on population vectors of neuron firing rates across all time bins (1s bins). The correlation values were then averaged to assess differences between groups.

#### Decoding group identity using neural features

This analysis aimed to decode the group identity (i.e., control, susceptible, or resilient, or saline vs. CNO for DREADD data) on a single-mouse basis by analyzing the pre-task activity, where no behavioral labels were available. As described in the “ Analysis of embedding dimensionality “ section, the pre-task activity can be represented as a geometrical object in the firing space, with each axis representing the firing rate of a neuron and each point in the space representing the activity of the neuronal ensemble in each time bin. We sought features that characterized the representational object and were invariant to rotations and scaling transformations, or a subset of these transformations, ensuring shape invariance of the object. We included only mice with at least 5 neurons simultaneously recorded during the pre-task period. For each mouse, we computed the cumulative variance explained across the principal components (PCs) (see “Analysis of embedding dimensionality” section for more details). We considered the cumulative values of the first three PCs as features for decoding. Following the inference of hidden states and the clustering analysis, we calculated the proportion of clusters retained at different thresholds and extracted the values at five distinct thresholds (see “Agglomerative Clustering Analysis” section). Additionally, we computed the mean and standard deviation of the spike count as the last two features. All of the neural features were computed using 1s bins to optimize the final decoding performance. Overall, we assessed a total of 10 neural features for each mouse.

We used the same Mahalanobis binary decoder procedure as previously described in the “Decoding group identity using behavioral features” section. Specifically in this case, the input to the binary classifier consisted of an *N* x *F* training matrix and a 1 x *F* testing matrix, where N represented the total number of training mice between the two classes, and *F* = 10 represented the total number of features. Prior to running the classification algorithm, we preprocessed the input matrices by applying a Min-Max Scaler to the mean and standard deviation of the spike count, ensuring that all features were scaled between 0 and 1 (because the PC cumulative variance and fraction of HMM clusters are defined between 0 and 1 by construction).

The same decoder procedure was also applied during the Pre-reward and Post-reward periods of the task. For the decoding using vCA1 activity, the training set was defined as 20% of the total number of mice due to the initial larger sample size.

#### Population decoding

Similar to previously described method (*46*), a linear support vector machine (SVM) classifier was trained to classify patterns of activity into two discrete categories. Results are reported as the generalized performance of the decoder using cross-validation with 80/20 training/testing split. Patterns of activity are defined as the mean firing rate during 0.5 s non-overlapping time bins. Pseudo-population recordings were generated by combining all neurons within the same region and the same group. As it is well known that neural activity in previous trials could strongly influence activity in current trials (*81*), for all pseudo-population decoding analyses, we balanced the number of trials of each trial type by taking into account both the previous and current trial types. In other words, we have equal numbers of water/water, sucrose/sucrose, water/sucrose, and sucrose/water trials (previous/current trials, respectively). Only neurons with at least 8 trials per each of the 4 trial types were included.

To decode current reward, we combined equal numbers of water/water and sucrose/water trials for water trials, and similarly, equal numbers of sucrose/sucrose and water/sucrose trials for sucrose trials. To decode previous reward, we combined equal numbers of water/water and water/sucrose trials for water trials, and similarly, equal numbers of sucrose/water and sucrose/sucrose trials for sucrose trials. To decode intention (stay vs. switch), we combined equal numbers of sucrose/sucrose and water/water trials for stay trials, and similarly, equal numbers of sucrose/water and water/sucrose trials for switch trials.

As each group may have different number of cells and trials, we used subsampling procedures to randomly subsample cells (60 neurons for both BLA and vCA1), and within those cells, randomly subsample trials equal to the group with the smallest number of trials. The resulting dataset was used to train SVM and obtain cross-validated decoding accuracies. For each set of subsampled cells, decoding accuracies across random subsampling of trials (repeated 10 times) were averaged to obtain a single sample of decoding accuracy. We repeated the whole procedure 10 times to obtain statistical comparisons across groups and against shuffled distribution.

For within-time-bin decoding, SVMs were trained using data from one time bin and tested using held-out data from the same time bin. For cross-time-bin decoding, SVMs were trained using data from one time bin and tested using data from the other time bins.

For statistical comparisons, decoding accuracy during Pre-reward (-4 to -3s) and Post-reward (0 to 1s) periods was averaged. If the mean decoding accuracy in a group is significantly higher than 2 x STD of its respective mean shuffled distribution, we then performed additional between-group comparisons (2-way comparison: Mann-Whitney test; 3-way comparison: Kruskal-Wallis test followed by Dunn’s multiple comparisons test).

#### Decoding switch vs. stay using HMM states

In addition to using recorded firing rates during the 4s pre-reward window to decode switch vs. stay, we also trained separate decoders using the smoothed activity of the hidden states inferred by the HMMs. This approach uniquely allowed us to identify population hidden states within this time window, and specifically those states that may be intention-selective, which can then be artificially manipulated to assess their necessity in decoding. It is important to note that the training of the HMM was performed on concatenated trials, which includes the four 1s bins pre-reward across all trial types. We then rearranged the sequence of hidden states in each trial type a posteriori.

Once the parameters of the HMMs were inferred, the models could smooth the observed data by computing the mean observed activity under the posterior distribution of hidden states (*53*). For instance, given the observed activity vector *X* during a time bin of a trial pre-reward, the HMM inferred a 0.2 probability of being in state *S* = 1 and a 0.8 probability of staying in state *S* = 2. More precisely, *P(S = 1 | X) = 0.2*, and *P(S = 2 | X) = 0.8*. The smoothed observations used to train and test the linear decoder were calculated as *Y = 0.2μ_1_ + 0.8μ_2_*, where *μ_j_* represents the inferred mean for the observations in state *j*.

To ensure robustness, we randomly sampled 60 neurons from each mouse for 10 neuronal subsamples. We generated 1000 pseudo trials for each of the four trial types, resulting in a total of 4000 pseudo trials for the training and testing sets, separately. The input data to train and test the decoder consisted of the smoothed activity assigned to each time bin. We trained and tested a support vector machine (SVM) classifier with a linear kernel, similar to the approach used in the population decoding using original firing rates, to decode switch vs. stay. In each cross-validation iteration, we randomly selected 100 pseudo trials as the training set and 20 pseudo trials as the testing set, for a total of 10 cross-validations. The final decoder accuracy was computed as the average across neuronal subsamples and cross-validations.

To assess the significance of the decoding signal, we compared it to a chance level, defined as 2 x standard deviations (STD) around the theoretical mean of the distribution of accuracies obtained after 1000 shuffles of the labels.

#### Defining intention-selective states

We conducted a detailed analysis of the distribution of hidden states across trial types to identify intention-selective states. For each mouse, we computed the fraction of occurrence of each hidden state within the 4s bins pre-reward across all trials. This distribution was then normalized to the total number of trials multiplied by the number of bins. We assessed this normalized distribution separately for each trial type.

Consistent with the decoding results, we observed that certain states appeared exclusively in either the stay or switch trials, with no occurrences in the other trial types. To quantify the amount of information each state held for the choice value, we computed the Shannon entropy (*82*). Specifically, for a given state, we normalized its occurrence frequency in each trial type to the total number of trials. The entropy of each state for the choice value was calculated using the following formula:

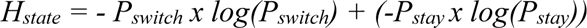

where *P_switch_* is the occurrence frequency of the state in switch trials (WS, SW) and *P_stay_ = 1 - P_switch_*. An entropy value of 0 indicates that the state provides highly informative signals for the intention. Therefore, we defined an intention-selective state as one with an entropy value of 0 for the choice value.

We examined the distribution of the fraction of intention-selective states at different clustering thresholds, and selected a threshold for each mouse that yielded the highest number of intention-selective states.

To examine the necessity and sufficiency of intention-selective states, we first excluded trials that contained intention-selective states in at least 3 time bins pre-reward. In the opposite approach, we enhanced the presence of intention-selective states in the decoding procedure by considering only those trials that included intention states in at least 3 time bins before the reward delivery.

#### Multi-dimensional scaling

To visualize the geometric structure of the data, we used Multi-Dimensional Scaling (MDS) transformation to obtain a low-dimensional representation of the data. For pre-task data, we started with the *N x F* matrix used for the Mahalanobis decoder, where N represents the total number of subjects across all three groups, and F denotes the number of features employed for decoding the group identity. Prior to the dimensionality reduction analysis, we normalized each group’s data by its variance to reduce noise and enhance the clarity of the final visualization.

Next, we performed a diagonalization of the dissimilarity matrix *N x N*, which contained the Euclidean distances between each pair of subjects in the feature space. We used the same procedure for the task period. In these cases, the input matrix was a *T x N* matrix, where *T* represents the total number of pseudo trials, and *N* denotes the number of neurons.

#### Inter-regional correlation

Firing rates (10ms bins) were averaged across all simultaneously recorded neurons in each mouse within the same region (BLA and vCA1). Then Pearson’s correlation was computed across simultaneously recorded regions within each 1s time window. The correlation was performed for each trial type separately, and a change in correlation (*corr_sucrose_ – corr_water_*) was computed as a measure to assess how different the inter-regional correlation is in sucrose vs. water trials for each animal.

#### Statistical analysis

No statistical tests were used to pre-determine sample size, but the sample sizes used are similar to those generally used within the field. Data were analyzed using parametric one- or two-way repeated measures (RM) ANOVA, or paired *t*-test. Where appropriate, ANOVA was followed by *post hoc* pairwise comparisons with corrections for multiple comparisons. If data were significantly non-normal (with *ɑ* = 0.05), non-parametric tests were used, including Kruskal-Wallis test or Mann-Whitney test (between-group comparisons) and Wilcoxon signed-rank test (within-group comparisons), and where appropriate, followed by *post hoc* comparisons with corrections for multiple comparisons. Categorical data were assessed using Fisher’s exact test. When comparing to chance, data was considered significant if it were outside of 2 x STD of chance distribution centered around theoretical chance level. Statistical comparisons between groups were performed for groups that were significantly different from respective chance distribution. All tests were two-sided. Statistical analyses were performed using Graphpad Prism V10.

---

**Fig. S1.**
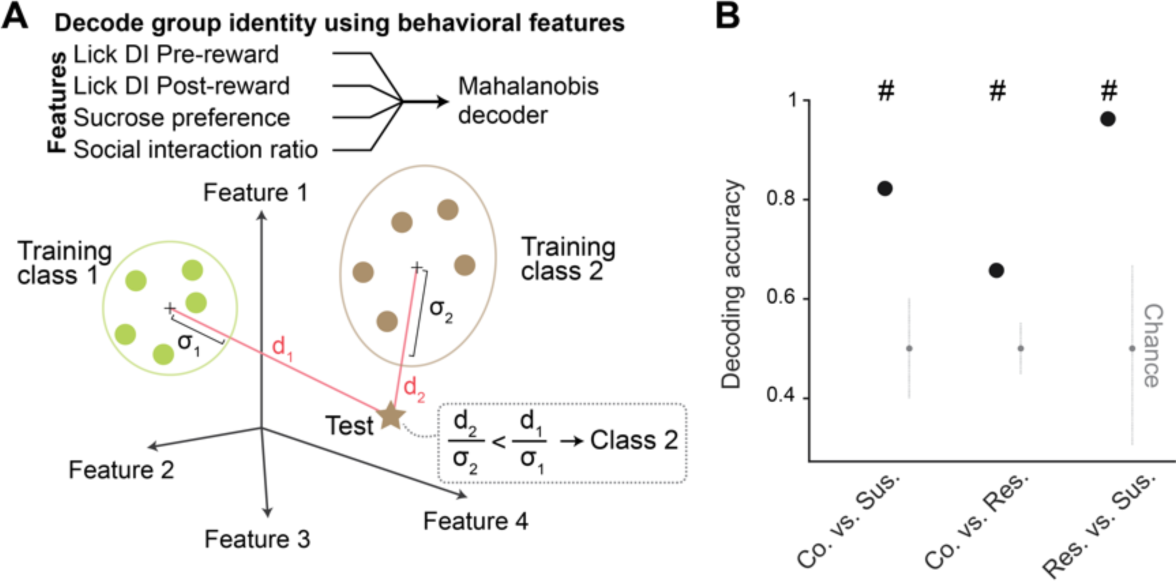
Decoding of group identity using behavioral features. (**A**) Schematic of the Mahalanobis decoder trained on behavioral features to decode group identity. (**B**) As further verification that behavioral features between groups classified using K-means clustering were different, group identity can be successfully decoded using Mahalanobis decoder trained on behavioral features including lick rate discrimination index (DI) during Pre- and Post-reward, sucrose preference, and social interaction ratio.

**Fig. S2.**
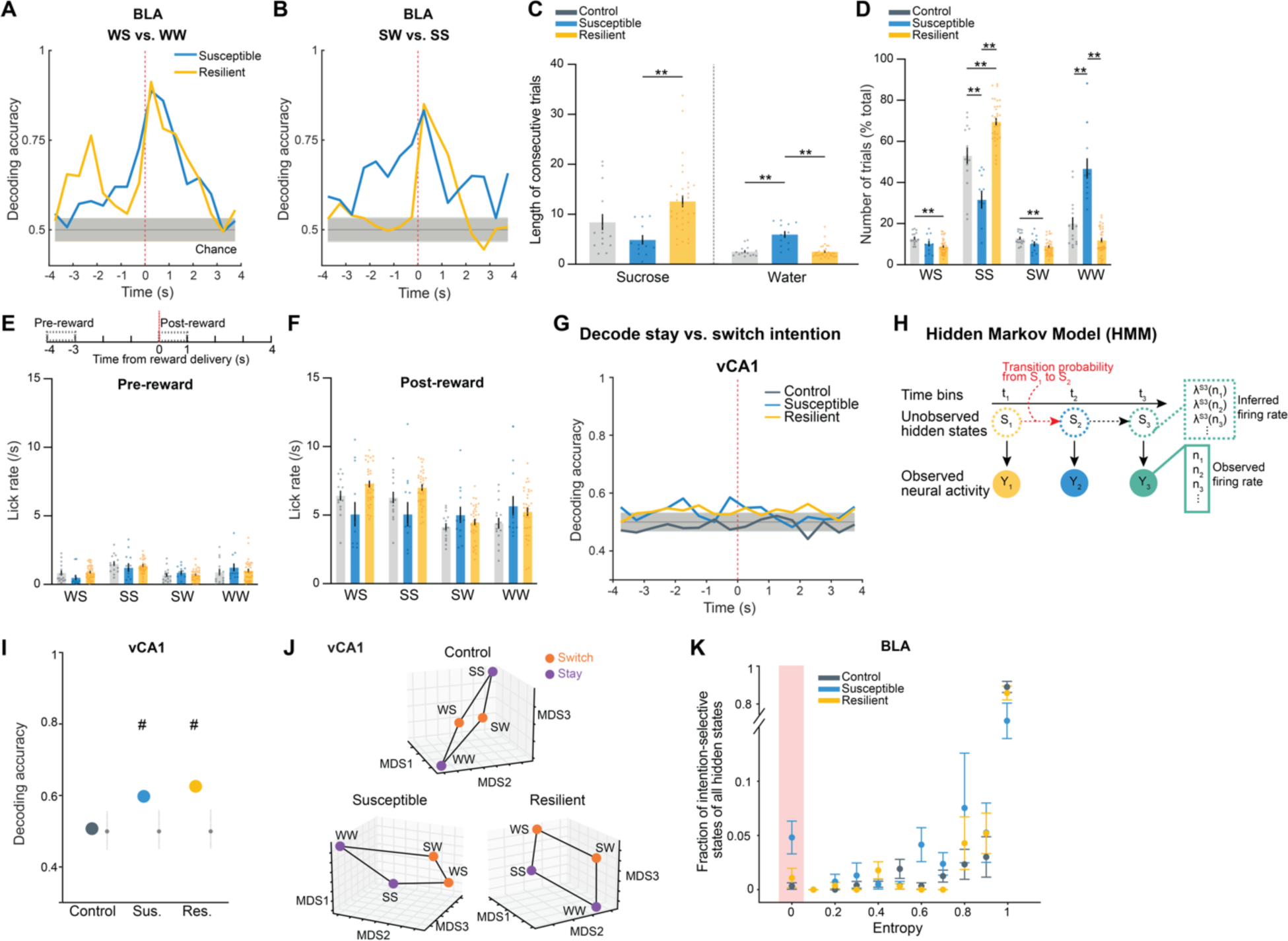
Intention selectivity in BLA as a unique susceptibility signature. **(A** and **B)** Decoding accuracy of reward choice in the current trial in the BLA when the previous trial was **(A)** water (WS vs. WW) or **(B)** sucrose (SW vs. SS). In resilient mice, upcoming reward choice can be decoded during Pre-reward time window when the previous trial was water, but not when previous trial was sucrose. In susceptible mice, upcoming reward choice during Pre-reward can be decoded regardless of the previous trial choice. **(C)** Susceptible mice showed fewer consecutive sucrose trials (ANOVA, effect of group: *F*_2,57_ = 7.60, *P* = 0.0012), and greater number of consecutive water trials in comparison to control and resilient mice (ANOVA, effect of group: *F*_2,57_ = 25.09, *P* < 0.0001). **(D)** Sucrose and water trials were further divided into sucrose/sucrose (SS), water/sucrose (WS), water/water (WW), and sucrose/water (SW) trials after taking into account the previous trial. Comparison of the proportion of trials in each of the 4 trial types revealed that controls showed greater proportion of switch trials (WS, SW). Resilient mice showed greatest proportion of SS trials, while susceptible mice showed greatest proportion of WW trials (RM-ANOVA, trial type x group interaction: *F*_6,171_ = 39.99, *P* < 0.0001). (**E** and **F**) Lick rates for each of the 4 trial types during (**E**) Pre-reward (RM-ANOVA, trial type x group interaction: *F*_6,171_ = 2.38, *P* = 0.031) and (**F**) Post-reward period (RM-ANOVA, trial type x group interaction: *F*_6,171_ = 9.80, *P* < 0.0001). **(G)** In vCA1, decoding accuracy of switch vs. stay intention using raw firing rates in susceptible and resilient mice was above chance. **(H)** Schematic of HMM to obtain population hidden states. **(I)** Similarly, in vCA1, decoding accuracy of switch vs. stay intention using inferred firing rates from HMM in the 4s preceding reward delivery in susceptible and resilient mice was above chance. **(J)** MDS visualization of inferred firing rates showed that population representations of switch and stay trials can be linearly separated in vCA1 neurons in susceptible and resilient mice than in controls. **(K)** Average distribution of the of fraction of intention-states across mice at different entropy values. For each mouse, the state entropy was computed at fixed threshold on the clustering dendrogram (see methods). Data are mean ± STD. Data are mean ± SEM unless otherwise stated. # Significantly different from chance; * *P* < 0.05, ** *P* < 0.01.

**Fig. S3.**
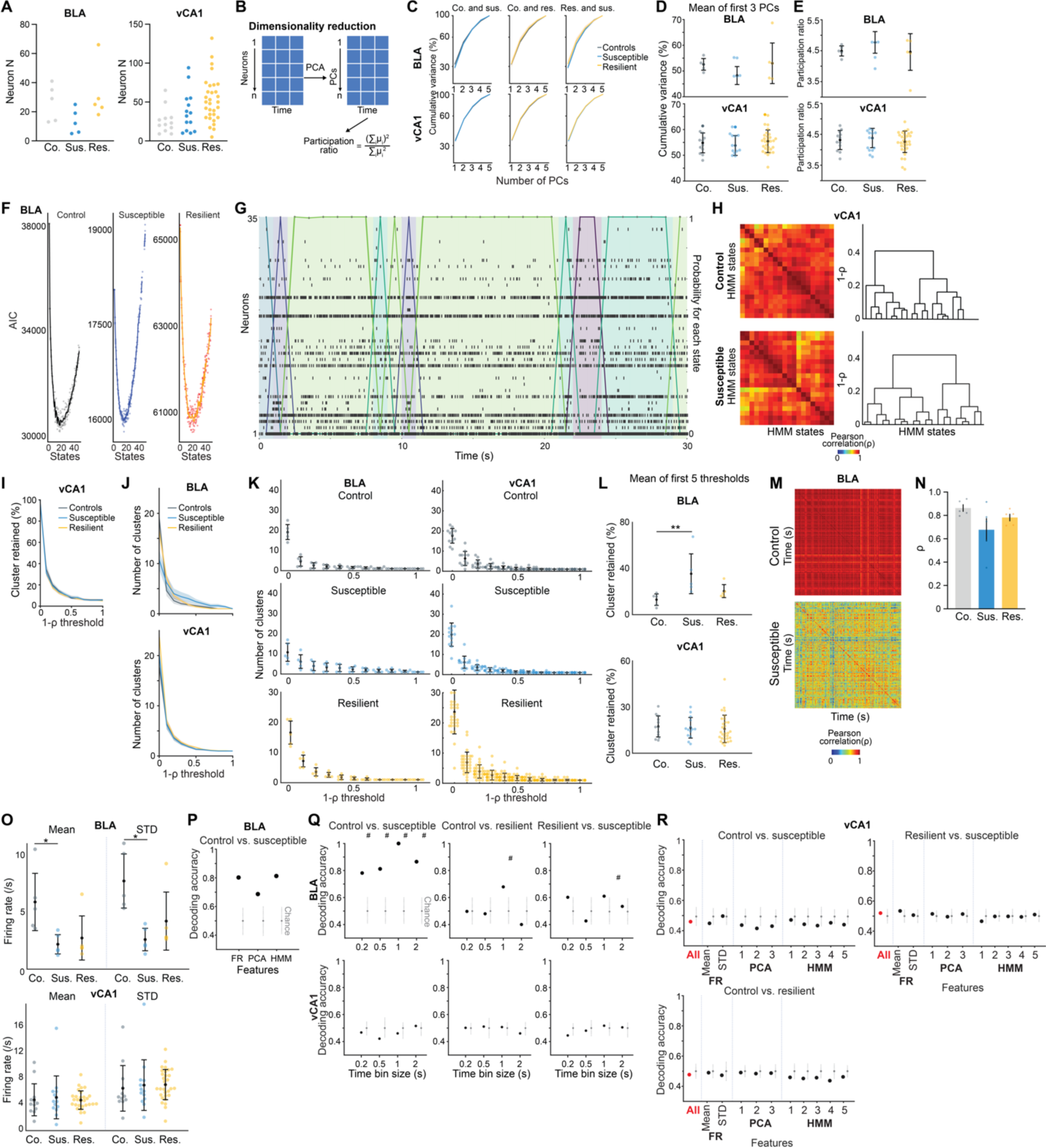
Distinct neural signatures of CSDS mice in the absence of task. (**A**) Number of neurons used in BLA and vCA1 in each mouse for analysis. (**B**) Schematic of dimensionality reduction using principal component analysis (PCA). The embedding dimensionality was quantified using participation ratio. (**C**) There were no statistically significant differences in cumulative variance explained by principal components (PCs) in BLA and vCA1 between groups. Data are mean ± SEM. (**D**) Mean of the cumulative variance of the first 3 principal components in BLA and vCA1. (**E**) Participation ratio of BLA and vCA1. (**F**) Akaike information criteria (AIC) from one example mouse in each group. HMM with the lowest AIC was selected as the best model. **(G)** Example raster and HMM states from one representative mouse. Colored lines indicate different hidden states and their posterior probability. (**H**) Two examples of HMM states correlation matrices for one control and one susceptible mouse, with respective dendrograms of agglomerative clustering in vCA1. **(I)** There was no difference in the proportion of distant hidden states in vCA1 between groups. **(J)** The number of clusters across thresholds did not differ between groups in BLA and vCA1. Data are mean ± SEM. **(K)** The number of clusters of individual mice. **(L)** Mean of the proportion of clusters in the first 5 thresholds showed that susceptible mice in BLA had greater proportion of unique hidden states in comparison to controls (Kruskal-Wallis, *P* = 0.0018). No group difference was found in vCA1. **(M)** Two example heatmaps of population activity correlation in BLA over time, showing that population activity patterns were much more correlated in the control (top) than the susceptible (bottom) mouse. **(N)** Average correlation of population activity across time in the BLA showed a trend towards lower correlated activity in the susceptible mice. **(O)** In BLA, susceptible mice showed reduced firing rates mean (Kruskal-Wallis, *P* = 0.020) and STD (Kruskal-Wallis, *P* = 0.0077) in comparison to controls. **(P)** Firing rate (FR), PCA, and HMM features each alone could successfully decode control vs. susceptible mice in BLA. **(Q)** Different time bin sizes were tested and the one that allowed the highest decoding accuracy between groups was chosen as the optimal bin size. **(R)** Group identity could not be decoded using Mahalanobis decoder trained on neural features in vCA1. The importance of each neural feature in decoding was examined by systematic removal of each of the features (subsequent columns). Data are mean ± STD, unless otherwise stated. Chance distributions are ± 2 x STD around theoretical chance level. # Significantly different from chance; * *P* < 0.05; ** *P* < 0.01.

**Fig. S4.**
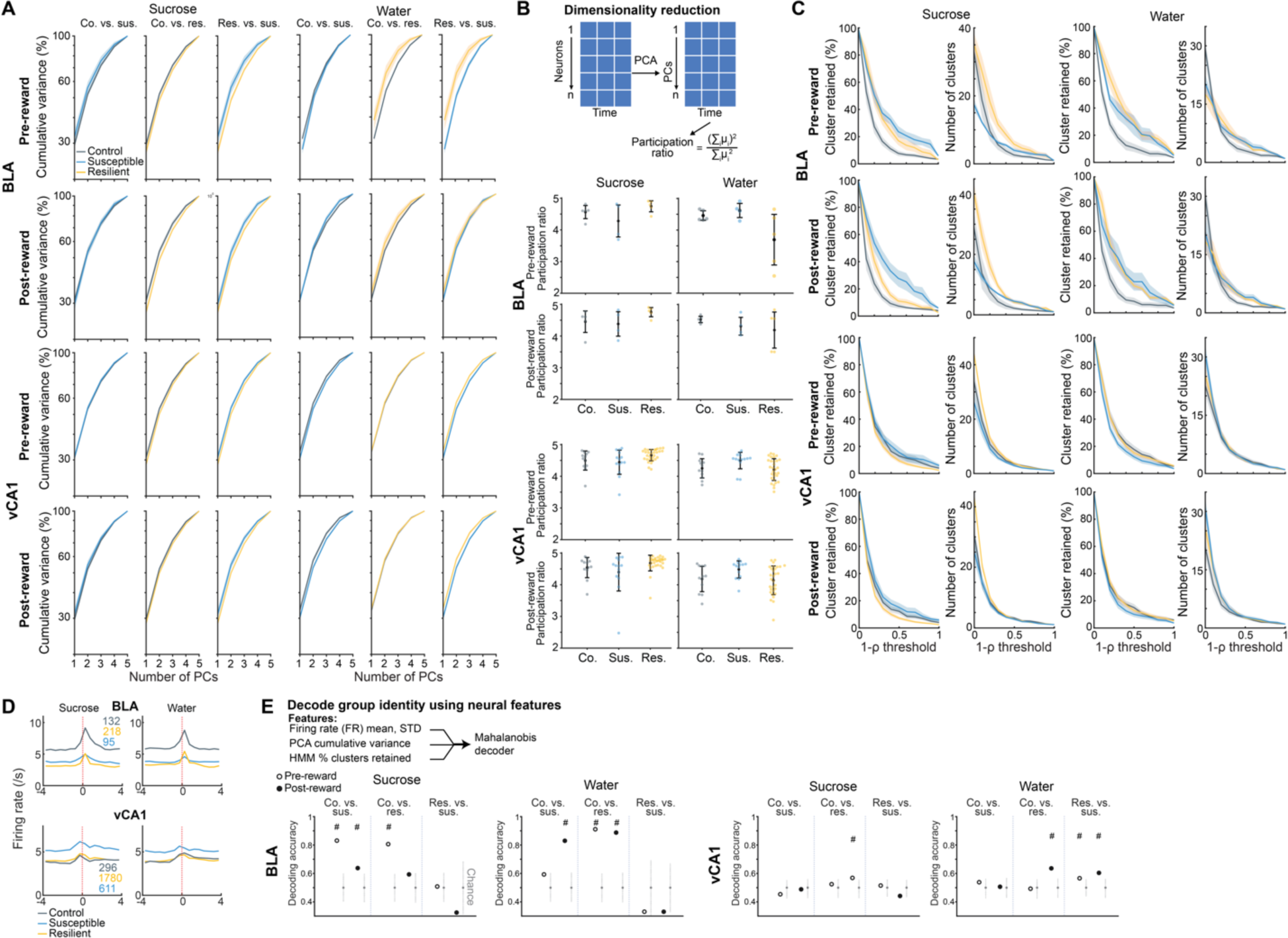
Dysfunctional single cell and population-level correlates for reward choice in mice susceptible to chronic stress. (**A**) Cumulative variance of PCs in BLA and vCA1 during Pre-reward and Post-reward periods revealed no difference between groups. (**B**) Participation ratio of BLA and vCA1 during Pre-reward and Post-reward periods showed no difference between groups. Data are mean ± STD. (**C**) The proportion and number of HMM clusters across different thresholds showed no statistical difference between groups, but BLA neurons in susceptible mice showed a trend towards higher proportion of unique clusters. (**D**) Trial-averaged firing rates of pseudo-populations of BLA and vCA1 neurons across groups. (**E**) Group identity could be decoded better than chance during specific time windows and trial types. Data are mean ± SEM unless otherwise stated. Chance distributions are ± 2 x STD around theoretical chance level. # Significantly different from chance.

**Fig. S5.**
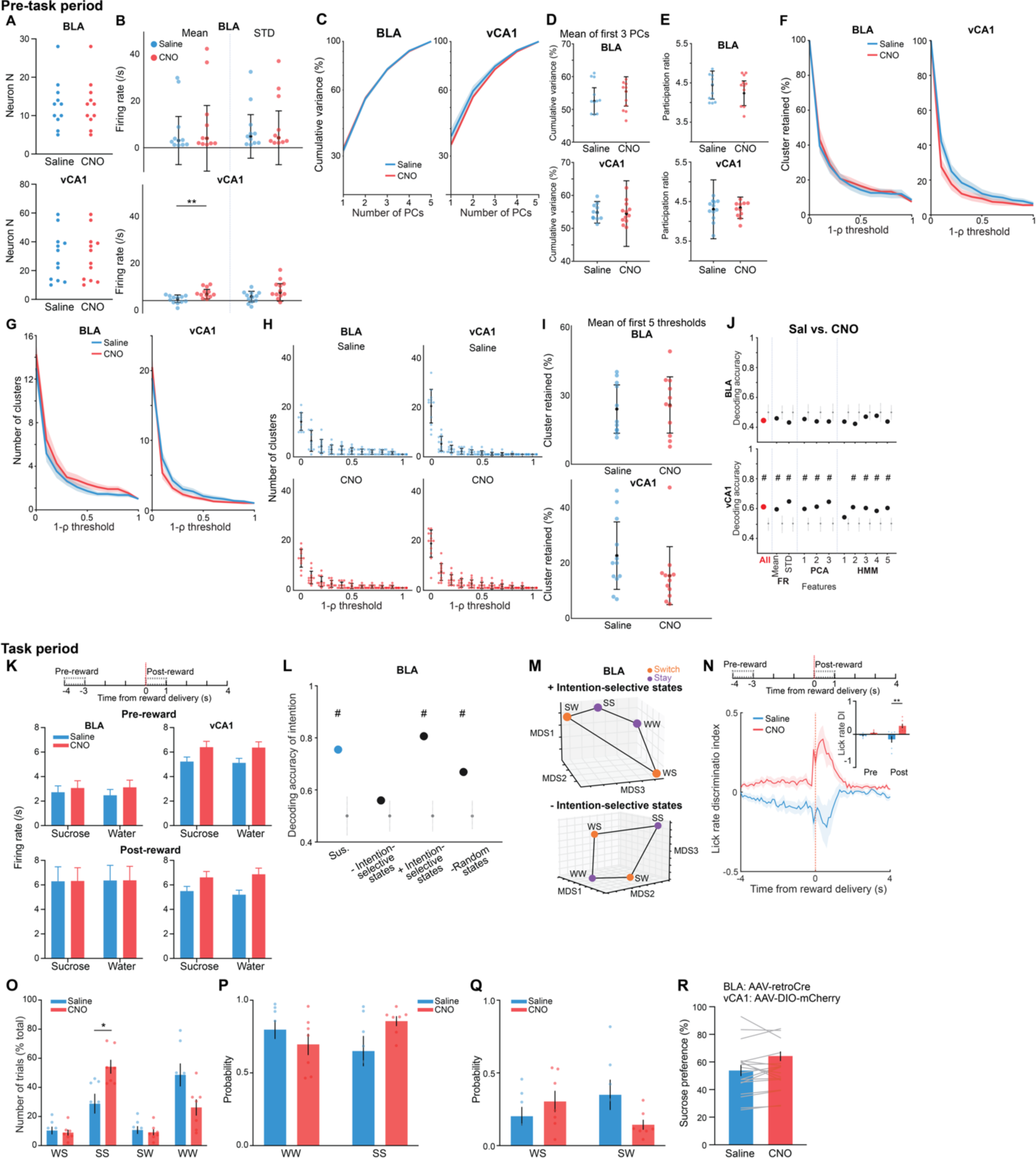
Rescue of dysfunctional vCA1-BLA activity and anhedonia by circuit-specific manipulations. (**A**) Number of neurons used in BLA and vCA1 in each mouse during saline and CNO period for analysis. (**B**) Mean vCA1 firing rates during pre-task period was increased following CNO (Mann-Whitney, *P* = 0.008). Data are mean ± STD. (**C**) Cumulative variance of PCs did not differ between saline and CNO periods in BLA or vCA1. (**D**) Mean of the cumulative variance of the first 3 PCs in BLA and vCA1. Data are median ± STD. (**E**) Participation ratio of BLA and vCA1. (**F**) Proportion of HMM clusters across different thresholds in BLA and vCA1. (**G**) The number of HMM clusters across different thresholds in BLA and vCA1. (**H**) The number of HMM clusters of individual mice across different thresholds in BLA and vCA1. Data are mean ± STD. (**I**) Mean of the proportion of clusters in the first 5 thresholds. Data are mean ± STD. (**J**) Despite no statistical difference in each of the FR, PCA, and HMM features, the decoder trained using all features could successfully decode between saline vs. CNO periods in vCA1 better than chance (± 2 x STD). (**K**) Firing rates of pseudo-population of BLA and vCA1 neurons during task showed that vCA1 neurons had elevated firing rates after CNO during both Pre-reward (RM ANOVA, effect of CNO: *F*_1,1092_ = 7.60, *P* = 0.0060) and Post-reward periods (RM-ANOVA, effect of CNO: *F*_1,1092_ = 9.57, *P* = 0.0020). **(L)** Removal of trials containing intention-selective states (-Intention-selective states) during the saline period reduced decoding accuracy of switch vs. stay trials to chance, whereas keeping only trials containing intention-selective states (+Intention-selective states) allowed successful decoding of stay vs. switch trials. Removal of trials with random states had little effect on decoding accuracy. Chance distributions are ± 2 x STD around theoretical chance level. **(M)** MDS visualization showed that keeping only intention-selective states allowed the representations of switch trials to be linearly distinguished from stay trials, whereas removal of intention-selective states prevents the representations of the two trial types from being linearly separated. **(N)** CNO increased lick rate discrimination index during Post-reward period in comparison to saline (RM-ANOVA, treatment x time interaction: *F_1,12_* = 10.80, *P* = 0.0065). **(O)** CNO increased the proportion of SS trials (RM ANOVA with Bonferroni’s multiple comparisons test, trial type x treatment interaction: *F*_3,36_ = 6.23, *P* = 0.0016). **(P)** CNO altered the proportion of stay (water-water and sucrose-sucrose) trials (RM ANOVA with Holm-Sidak’s multiple comparisons test, effect of group: *F*_1,12_ = 5.61, *P* = 0.036). **(Q)** CNO altered the proportion of switch (water-sucrose and sucrose-water) trials (RM-ANOVA, effect of group: *F*_1,12_ = 5.61, *P* = 0.036). **(R)** Mice microinfused with the control virus (AAV-DIO-mCherry) and given CNO showed no change in sucrose preference. Data are mean ± SEM unless otherwise stated. # Significantly different from chance; * *P* < 0.05, ** *P* < 0.01.

